# Differential expression of H2A isoforms contribute to tissue and lineage specificity with HIST2H2AC as a potential cancer biomarker

**DOI:** 10.1101/2021.03.10.434738

**Authors:** Sanket Girish Shah, Mudasir Rashid, Abhiram Natu, Sanjay Gupta

## Abstract

Recent advancements in the field of histone biology imply non-redundancy in the function of histone H2A isoforms; however, the expression of H2A isoforms in various normal tissue types, the correlation among organs and tumor/tumor type-specific expression remain poorly investigated. The profiling of sixteen H2A isoforms in eleven different normal human tissue types strongly suggests their tissue-specific or predominant expression. Further, clustering analysis shows a lineage-specific correlation of H2A isoforms. In continuation, the expression analysis in twelve human tumor types shows overexpression of HIST2H2AC. Moreover, overexpression was observed exclusively in tumor samples but not with fetal samples; highlighting the cancer-specific association of HIST2H2AC. Further, *in silico* analysis of TCGA pan-cancer data also showed tumor-specific over-expression of the HIST2H2AC isoform. Our findings provide insights into tissue-type-specificity of histone H2A isoforms expression patterns and advance our understanding of their importance in lineage specification and cancer.

## Introduction

Distinct tissue-specific gene expression patterns are usually formed during early embryonic development in an organism which is crucial for cellular differentiation and organ formation (Yi et al., 2010). These expression alterations are often associated with changes in the epigenetic landscape. The key regulators of epigenome such as DNA methylation, incorporation of histone variants, and histone post translational modifications have been extensively studied. However, emerging shreds of evidence suggest that the replication-dependent histone isoforms may constitute a functional layer of chromatin regulation (Jones and Martienssen, 2005). Histone isoforms, unlike histone variants, are organized in clusters, transcribed during the S phase of the cell cycle, and vary by a few amino acids at the protein level thus making them disparate from histone variants at the organizational, transcriptional, and protein level (Singh et al., 2013). Further, due to their high degree of similarity among protein sequences, unavailability of antibodies, and a presumption that they are functionally redundant, the importance of histone isoforms in providing epigenetic plasticity is not well studied.

Recently, our study has shown advances in the complexity of eukaryotic H2A isoforms, their roles in defining nucleosome organization, and their association with diseases (Shah et al., 2020). H2A isoform family consists of 16 genes and codes for 11 proteins (Singh et al., 2013). However, reports suggest that H2A isoforms could be functionally non-overlapping. The knockdown of the HIST1H2AC isoform leads to an increase in cellular proliferation and tumorigenic potential in the bladder cancer cell line (Singh et al., 2013). Contrary to an earlier report, knockdown of the same isoform leads to loss of cellular proliferation, G0/G1 cell cycle arrest, and defective estrogen signaling in breast cancer cells (Su et al., 2014). The level of HIST1H2AC isoform was also found to be elevated in chronic myeloid leukemia patients (Singh et al., 2015; Olivares et al., 2009; Singh et al., 2007). Further, the homolog of H2A1C in rat, H2A.1 was also overexpressed in hepatocellular carcinoma (Bhattacharya et al., 2018). Another isoform, HIST2H2AC was found to play an oncogenic role, downstream of EGFR signaling in breast cancer cells (Monteiro et al., 2017). These studies suggest that the altered levels of H2A isoform associates with pathophysiological states.

Comprehensive profiling of H2A isoforms in normal, undifferentiated, and different cancer types has been carried out to highlight their importance in tissue-specificity and potential importance in cancer. Our study highlights the tissue and lineage-specific expression of H2A isoforms. Also, tumor-specific expression of HIST2H2AC in different tumor types strongly supports that this H2A isoform can be exploited as early predictive biomarkers for cancer.

## Results

### Predominant and specific expression of H2A isoforms in different normal tissues

H2A isoforms were profiled and compared across eleven different normal tissue types and they were found to be expressed in tissue-specific and/or predominant (Figure1, Figure1-figure supplement1). HIST1H2AA showed tissue-specific expression as expressed ~1000X higher in testis compared to other organs. Further, other isoforms were differentially enriched in different tissues with the fold change of 2 to 15 suggesting their predominant expression in particular tissues. The genes like HIST1H2AB/HIST1H2AE and HIST1H2AG/HIST1H2AI/HIST1H2AK/HIST1H2AL /HIST1H2AM which code for the same proteins have shown similar expression profile in tissue predominant manner. However, in addition to major similarity in expression profile in all tissues, HIST1H2AB showed enrichment in testis (~2X); whereas, HIST1H2AE was upregulated in the breast tissue. Further, among HIST1H2AG/I/K/L/M, HIST1H2AL has increased expression in the tongue; whereas, HIST1H2AM was increased in the rectum and colon.

**Figure 1:**
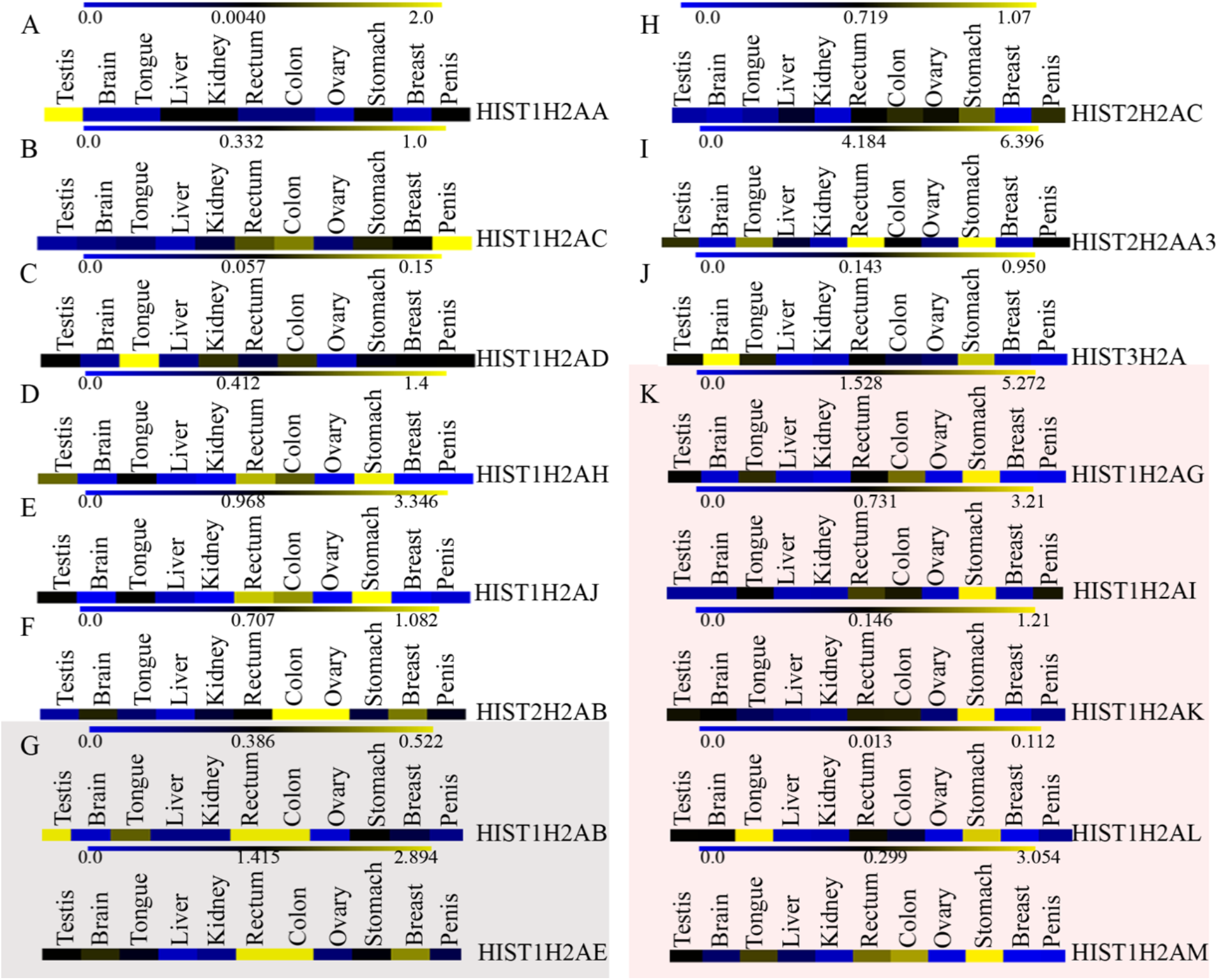
Tissue-specific and predominant expression of histone H2A isoforms in eleven different normal tissue types: The data for all H2A genes were normalized to the expression levels of total H2A as a common reference point. (A-K) Heatmap representation of H2A isoforms coded either by a single gene (white color) or two genes (grey color) or five genes (light pink color). The blue-black-yellow color scale displays low, median, and high expressions, respectively. (**A)** HIST1H2AA shows testis specific expression (~1000 fold higher expression than median value); **(B & C)** HIST1H2AC and HIST1H2AD are predominantly expressed in penis (~4 fold upregulation than median value) and tongue respectively (~3 fold higher expression than median value); (**D & E)** High expression of HIST1H2AH and HIST1H2AJ in both stomach and rectum tissues (~4 fold for HIST1H2AH and ~3 fold for HIST1H2AJ, upregulation compared to median value); (**F)** Increased expression of HIST2H2AB in colon and ovary (~2 fold higher expression than median value); (**G)** HIST1H2AB and HIST1H2AE isoforms code for a same proteins and both have high expression in rectum and colon (~2 fold upregulation compared to median value); (**H)** HIST2H2AC does not have predominant expression in any tissue type; **(I & J)** High expression of HIST2H2AA3 in stomach and rectum (~2 fold) and of HIST3H2A in brain (~5 fold) (**K)** All five genes, HIST1H2AG, HIST1H2AI, HIST1H2AK, HIST1H2AL and HIST1H2AM code for the same protein and all are predominantly expressed in stomach (~5-10 fold higher expression than median value).

To further identify the correlation between the expression of H2A isoforms and cell lineage of different organs, hierarchical clustering analysis was carried out. The analysis showed that the different organs form clusters according to their respective cell lineages. Brain and breast tissues form one cluster, which belongs to the ectoderm lineage. The tongue, rectum, colon, and stomach forms one family which belongs to the endoderm lineage, whereas the kidney, liver, penis, and ovary are formed under one tree, which is mesoderm lineage (Figure 2A). Though the liver is mainly derived from endoderm, it comes under the tree of mesoderm as the liver contains both origin cell types i.e. hepatoblast, kupffer cells, and hepatic mesenchyme cells. Surprisingly, testis did not fall into any family. This might be due to testis-specific expression of HIST1H2AA, and absence of other isoforms.

**Figure 2:**
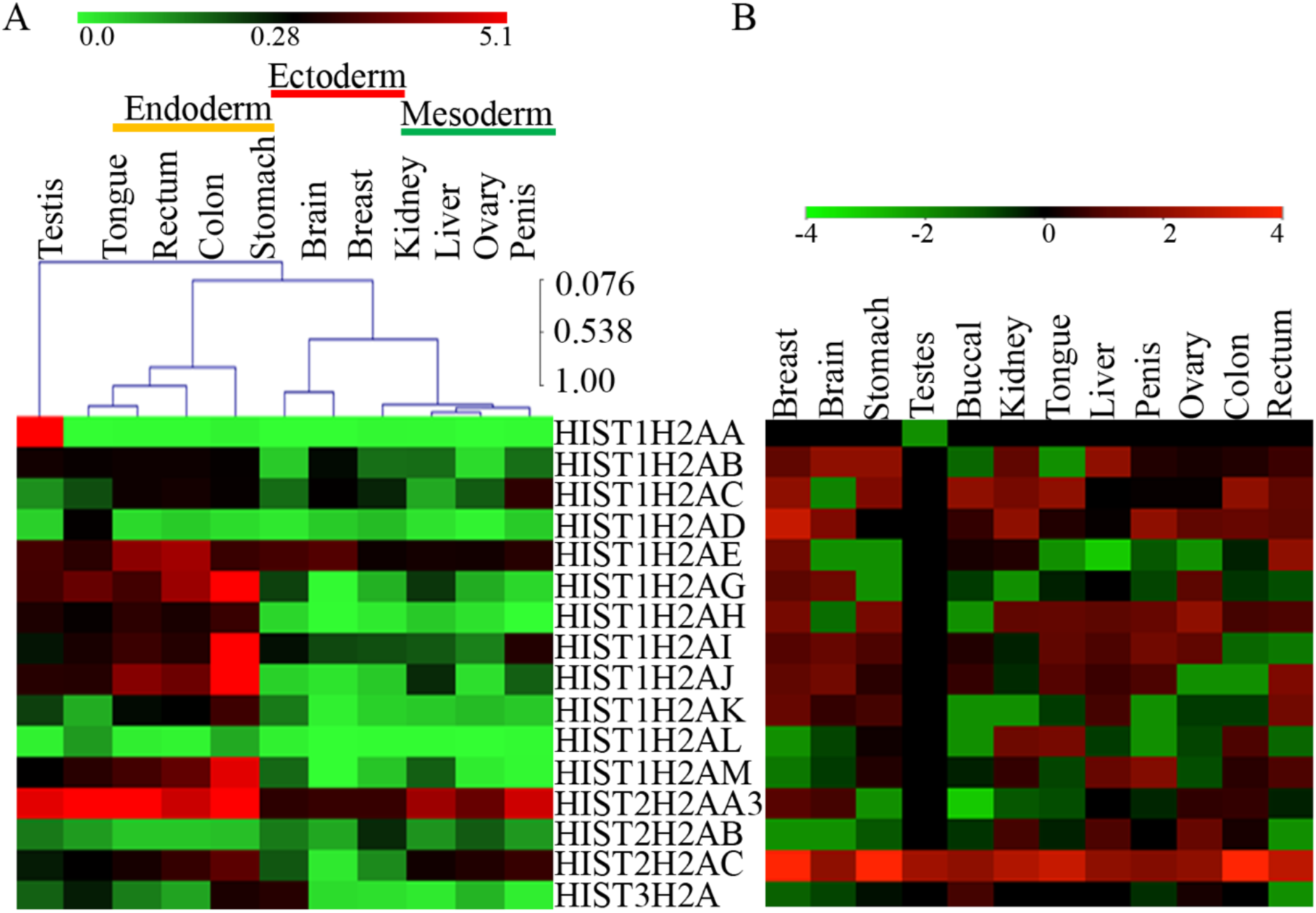
Lineage-specific correlation of H2A isoforms and altered expression profile in various tumors: (**A)** Heatmap representation with clustering analysis for all normal tissues and all H2A isoforms. The data for expression value is normalized with reference gene and plotted as a heatmap with the green-black-red color scheme. Hierarchical clustering analysis with complete linkage shows the lineage-specific expression of H2A isoforms. Three lines above the heatmap represent organs specific for ectoderm, mesoderm, and endoderm. Testis does not fit into any lineage due to the 1000-fold upregulation of HIST1H2AA in it. (**B)** Heatmap representation with green-black-red scale for all sixteen H2A genes in twelve different tumor types. The mean expression of each gene for individual tumor type is calculated from individual fold change values of each sample and represented here.

Taken together, data suggests tissue-specific/predominant and lineage specific expression of H2A isoforms. Further, all H2A isoforms were profiled in different origins of tumor tissue samples (Figure2B, Figure2B-figure supplement1).

### Tumor-specific expression of HIST2H2AC

HIST2H2AC was found to be universally upregulated in all of the tumor types along with the differential expression of other isoforms in diverse cancers (Figure2B & Figure3). Further, the expression of HIST2H2AC was analyzed in the TCGA PANCAN cohort. The median expression of HIST2H2AC was high in the tumor samples data set (n=9078) compared to the normal samples set (n=677) (Figure3-figure supplement 1A). Further, the stage-wise analysis showed that HIST2H2AC expression increases in stage I compared to normal samples and remained unchanged in later stages (Figure3-figure supplement 1B).

**Figure3:**
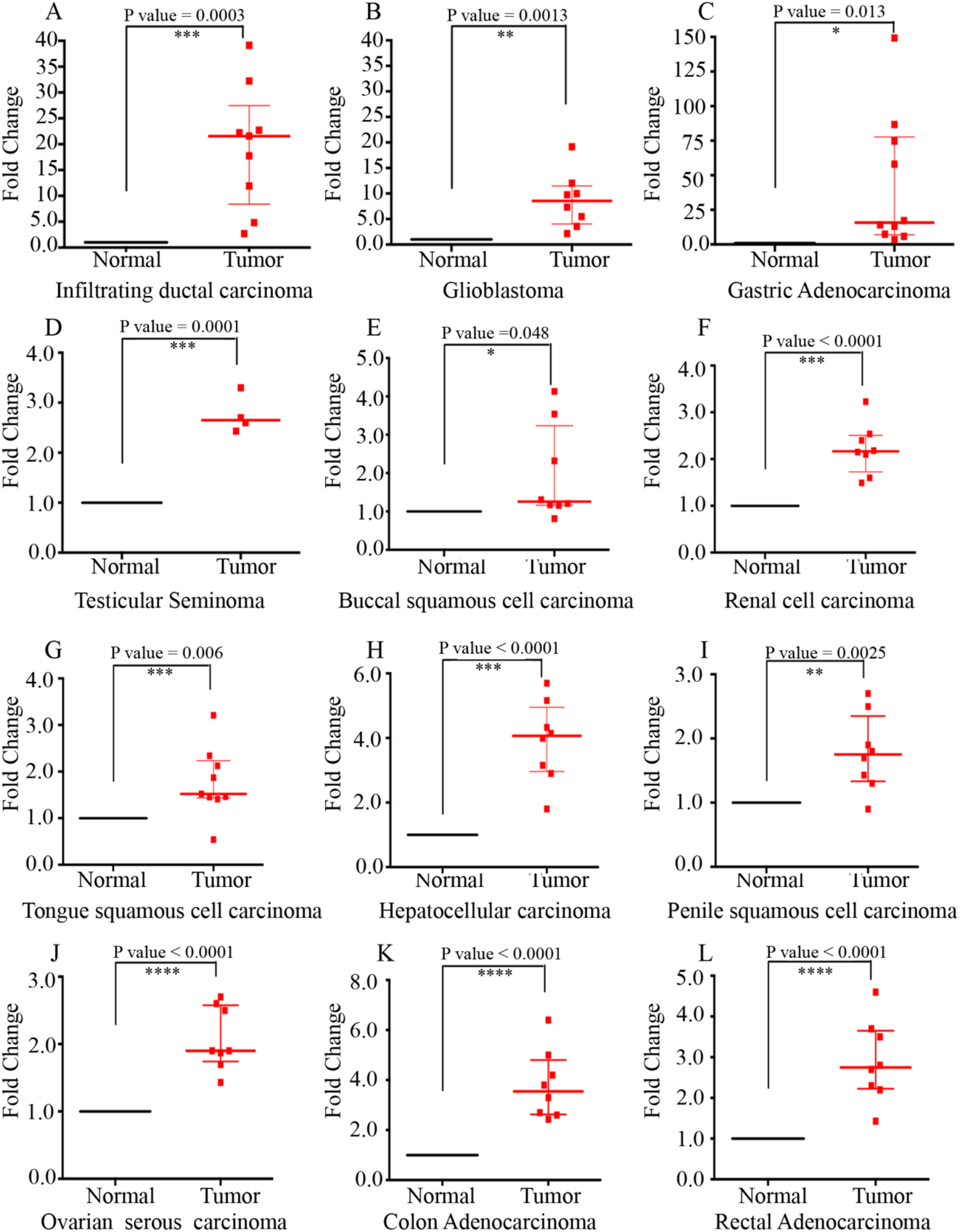
Overexpression of HIST2H2AC in multiple tumor tissue types: **A-L,** Scatter dot plot representation of HIST2H2AC expression in twelve different tumor types. The fold change for each sample is calculated with reference to the normal sample and represented here. Each dot represents a single sample, black line and the red line represents median expression value and error bars display 25 and 75 percentile values. p<0.05=*, p<0.01=** and p<0.001=***.

### Enhanced expression of HIST2H2AC expression is associated with dedifferentiated state

To validate whether the cancer-specific overexpression of HIST2H2AC is an oncofetal phenomenon, altered isoforms were profiled in the normal, tumor, and fetal RNA samples of the brain, kidney, and liver (Figure 4A, 4B and 4C). HIST2H2AC was overexpressed exclusively in all the tumor samples; whereas other isoforms coding for different proteins, HIST1H2AB/HIST1H2AD, HIST1H2AC/HIST1H2AD, and HIST1H2AB were upregulated in the tumor as well as fetal samples of brain, kidney, and liver, respectively (Figure 4A and B); whereas, HIST1H2AE is downregulated in tumor and fetal samples liver (Figure 4C). HIST1H2AM remained unaltered in all tissue samples and origins. In line with previous research on histone variants, these differentiation and dedifferentiation processes are not randomly defined. Tissue-specific expression of histone isoforms and their incorporation in the chromatin may contribute to a specific epigenetic landscape for defining an altered gene expression pattern in tumor phenotype (Figure 4D).

**Figure4:**
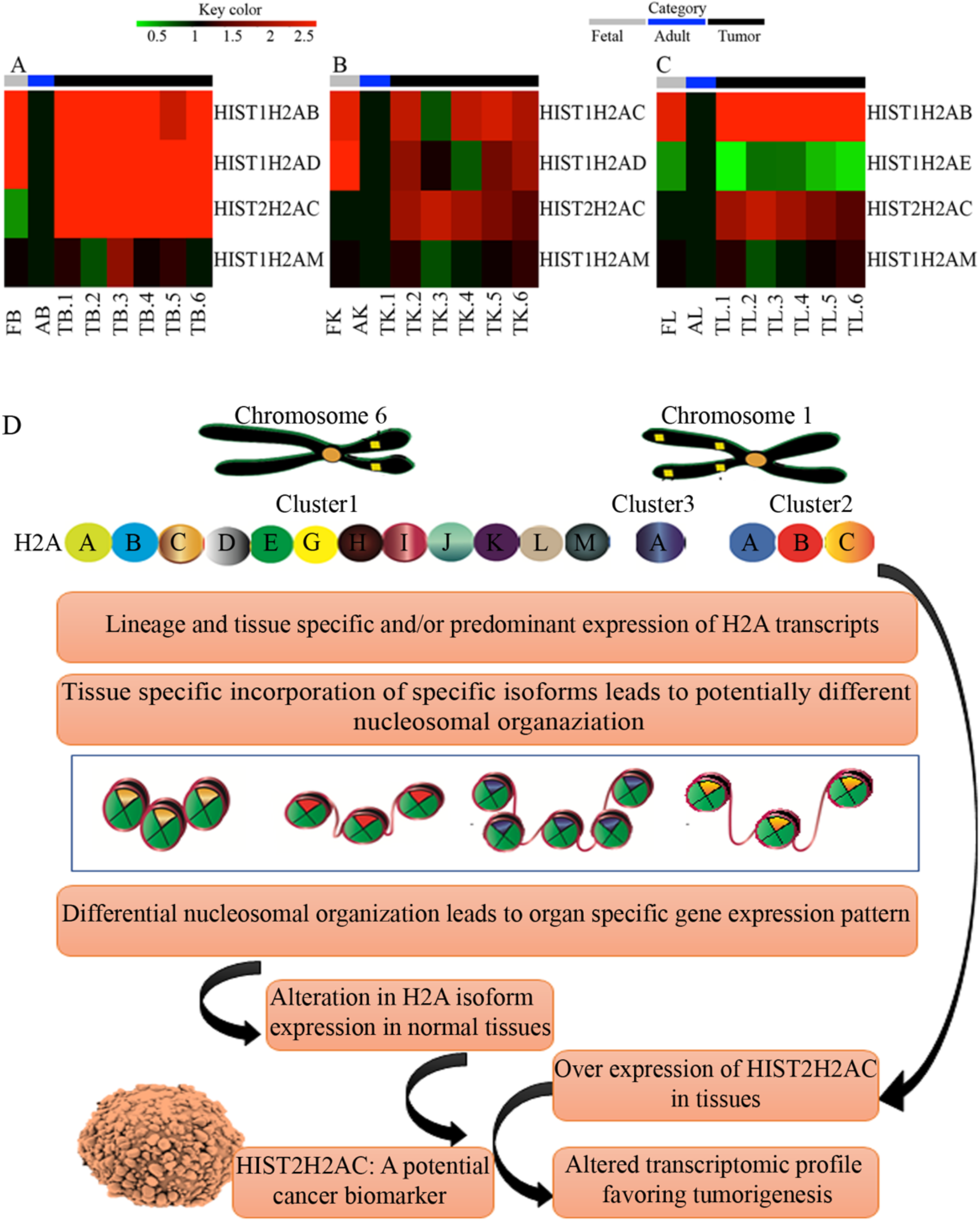
HIST2H2AC is associated with dedifferentiation phenotype and schematic representation highlighting the potential role of H2A isoforms. (**A-C)** Heatmap representation of fetal, normal, and tumor tissue samples for brain, kidney, and liver representing ectoderm, mesoderm, and endoderm, respectively. Altered isoforms in cancer for each category are taken further and represented along with normal and fetal tissues. Grey, blue and black color represents fetal, normal (adult), and tumor categories, respectively. HIST2H2AC shows high expression only in tumor tissues of all three origins but not in fetal tissues. The other isoforms show alteration both in tumor and fetal tissues compared to normal tissues. (D) Model for organization and expression of H2A isoforms. H2A isoforms are organized in three clusters. Cluster 1 is organized on chromosome 6; whereas, cluster 2 and cluster 3 are organized on chromosome 1. These isoforms are predominantly expressed in a lineage and tissue-specific manner. Differential incorporation of specific isoforms in specific tissues leads to a potentially diverse nucleosomal organization which eventually leads to organ-specific gene expression. The expression of H2A isoforms alters in cancer with an increased tumor-specific expression of HIST2H2AC. High expression of HIST2H2AC in tumor samples (n=96) of different origins highlights its utility as a potential cancer biomarker.

## Discussion

Earlier studies have shown that transcription factors play a critical role in shaping and regulating cellular identity (Peñalosa-Ruiz et al., 2019). Epigenetic factors like DNA methylation, non-coding RNAs, histone modifications, and histone variants contribute in defining the unique gene expression profile that gives rise to specific cellular identity in different physiological states (Fan et al., 2015; Filipescu et al., 2014). Our present study provides an interesting facet of tissue-specific and predominant expression of H2A isoforms along with the lineage-specific association. Exclusive expression with higher fold and low expression of other genes will determine tissue specificity (Fagerberg et al., 2014; Sonawane et al., 2017). The differentiated cell inherits specific gene signatures from the parental cell because of the epigenomic memory. Such a signature pattern does not include only origin-specific transcription factors but also the histone genes (Filipescu et al., 2014).

The tumor-specific increased expression of HIST2H2AC in multiple tumors might contribute to the tumor-specific chromatin organization for altered gene expression and tissue phenotype. Further validation in a bigger cohort and serum samples will strengthen its utility as a predictive molecular biomarker. Additionally, an understanding of how the expression of HIST2H2AC leads to distinct molecular profiles in tumors will shed light on the tumor-specific function and potential utility in cancer. HIST2H2AC is not predominately expressed in normal tissue samples. Unlike oncofetal genes, like alpha-fetoprotein, chronic embryonic antigen, HIST2H2AC was found to be associated exclusively with dedifferentiated tumor samples and not with the undifferentiated fetal tissue samples.

The overexpression of a specific H2A isoform in tissues or during cancer implies that the protein governs a global nucleosomal organization thereby altering transcription factor binding and transcription kinetics (Osakabe et al., 2018). Previously our lab has shown the molecular basis of H2A isoforms in regulating nucleosome stability to attain a physiological state (Bhattacharya et al., 2018). In continuation, the lab has shown *in silico* differential nucleosomal stability of different H2A isoforms that differ among themselves by few amino acids (Shah et al., 2020). Therefore, the tissue-specific expression and incorporation of H2A isoforms in genomic regions may provide specific nucleosomal organization for defining cell fate. Moreover, post-translational modifications may provide a second level of plasticity to maintain different transcriptional stages for gene expression. Such changes will increase cell-to-cell variability in gene expression and other phenotypes.

In conclusion, the genomic sequence along with histone isoforms and variants define a chromatin landscape to ensure proper gene expression and cell fate.

## Materials and Methods

### Human normal RNA

Total RNA for eleven different normal tissues was purchased from Agilent Company such as kidney (540013), rectum (540069), breast (540045), stomach (540037), ovary (540071), testis (540049), liver (540017), colon (540009), tongue (540149), penis (540147), and brain (540005). RNA for the fetal liver was obtained from Agilent (540173) and RNA for the fetal brain (636526) and kidney (636584) was procured from Takara. The obtained normal and fetal RNA was a pool of 4-5 samples for each tissue type.

### Collection of human patient tissue samples

Samples for twelve different human tumor tissue types were collected from ACTREC Biorepository (ACTREC-TTR) and TMH-INTTR and cryopreserved at −80°C. The project was approved by Institute human ethics committee vide #164 dated 27-04-2015. The inclusion and exclusion (naïve and no biological hazard) criteria for all the collected samples have strictly adhered. All collected tissue samples were grade III to IV. The detail of sample size and cancer type is provided in Figure2-figure supplement1; adenocarcinoma of stomach, colon, rectum, and ovary; infiltrating ductal carcinoma of the breast; squamous cell carcinoma of head and neck (tongue, buccal mucosa) and penis; hepatocellular carcinoma of the liver; glioblastoma of the brain; renal cell carcinoma of kidney and seminoma of testis were retrieved from EMR database of TMH (http://10.100.52.12/emrtmhact/default.asp). Histopathology analysis of tissues was confirmed earlier (Shah et al., 2019). The samples with more than 70% tumor content were processed for further studies.

### Total RNA quality assessment, quantitation, cDNA synthesis, and qPCR

The protocol was adapted from Shah et al 2019. In brief, total RNA was isolated from human normal and tumor tissues according to manufacture protocol (Agilent Cat#400800), and the quantity and quality were confirmed by NanoDrop 2000C spectrophotometer, and agarose formaldehyde denaturing gel electrophoresis. Synthesized cDNA was used as a template for a qRT-PCR reaction. Primers used for qRT-PCR reactions for all H2A isoforms and total H2A are provided in Figure1-figure supplement 1. The fold change was calculated directly from the ΔCt value with normalization from the reference gene (Total H2A) for normal RNA samples and represented as a heatmap. For tumor samples, normalization was done using ΔΔCt method, and fold change was calculated with reference to normal samples. Further, this fold change was log2 transformed and plotted as a heatmap to represent all tumor samples and all H2A genes together. For fetal and tumor tissue samples, fold change was normalized with normal (adult) samples and plotted as a heatmap.

### TCGA data analysis

HIST2H2AC expression was analyzed in pan-cancer cohort of normal and tumor samples. TCGA PANCAN normalized RSEM (log2 X+1) counts were obtained from the UCSC Xena browser. PANCAN analysis was carried out with the use of R3.3.3 software using the boxplot function and with the use of clinical parameters from cbioportal (http://www.R-project.org/).

### Heatmap representation and clustering analysis

MeV software was used for clustering analysis. The mean value of three independent experiments was taken and the heatmap was plotted using MeV. Using Hierarchical clustering and complete linkage, a sample tree was formed to find out the correlation among different tissue samples.

### Statistical analysis

Scatter dot plot representation was used to show fold change difference between normal and tumor samples (GraphPad Prism6.0). The mean value of technical and biological replicates of each of the samples was used for representation. The p-value for scatter dot plot was plotted using an unpaired t-test. The p-value for TCGA data was determined using Wilcoxon–Mann– Whitney test.

## Acknowledgment

SGS, MR, and AN are supported by the DAE-ACTREC fellowship. We are thankful to Dr. Poonam Gera, Officer-In-Charge, ACTREC-TTR, Ms. Manisha Kulkarni, and Mr. Anand Deshpande from TMH-INTTR for providing tissue samples.

## Additional information

### Author contribution

SGS and SG, Conception and design, Acquisition of data, Analysis and interpretation of data, Manuscript writing; MR, Acquisition of data, Analysis and interpretation of data; AN, Analysis and interpretation of data, manuscript writing.

### Ethics

The project IRG#164 was approved by the ethical committee of TMH (Tata Memorial Hospital).

## Supplementary Figures

**Figure1-figure supplement1:**
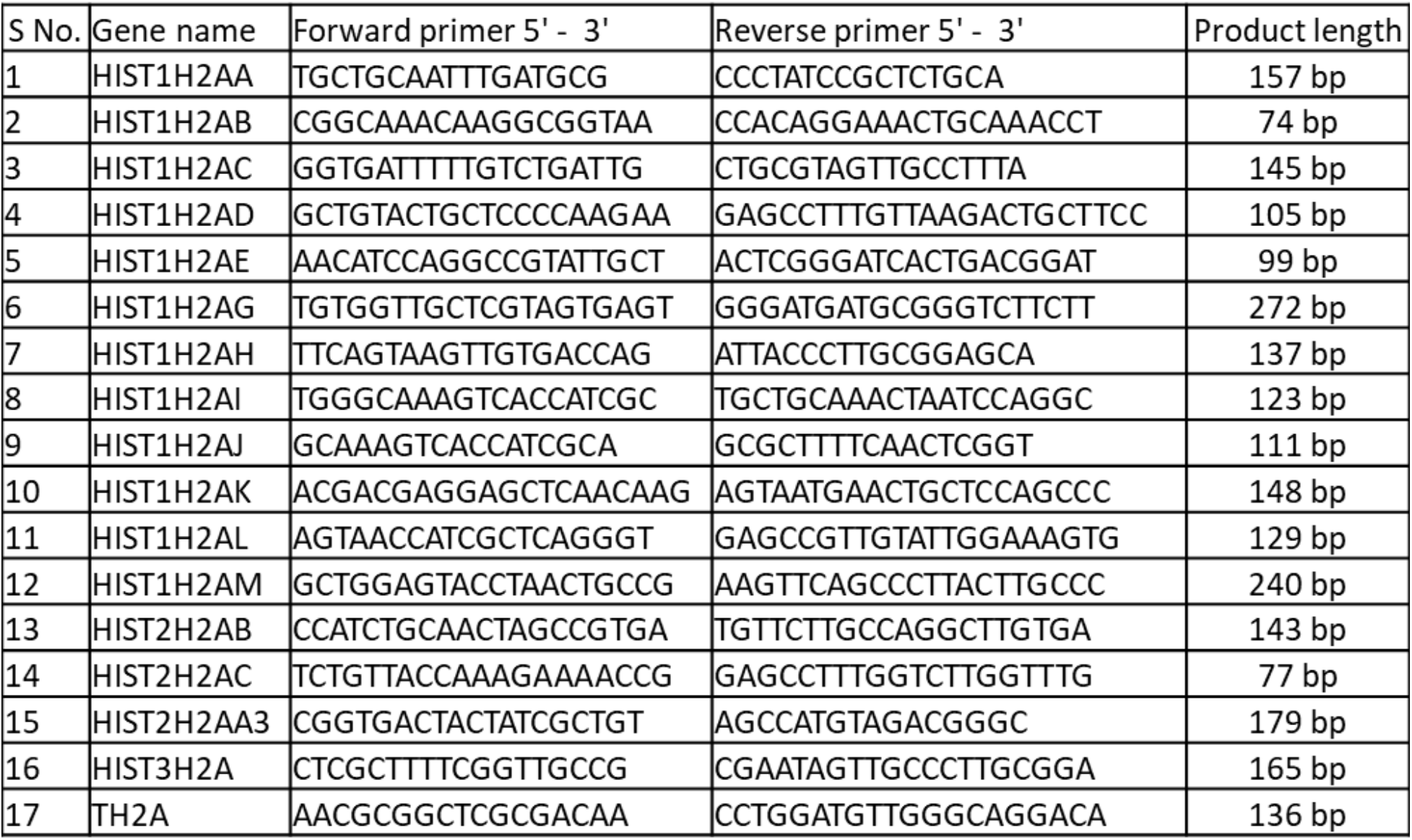
Details of primers of human H2A isoforms: Figure representing seventeen different set of forward and reverse primers along with their product size. Total H2A was used for normalization.

**Figure2-figure supplement1:**
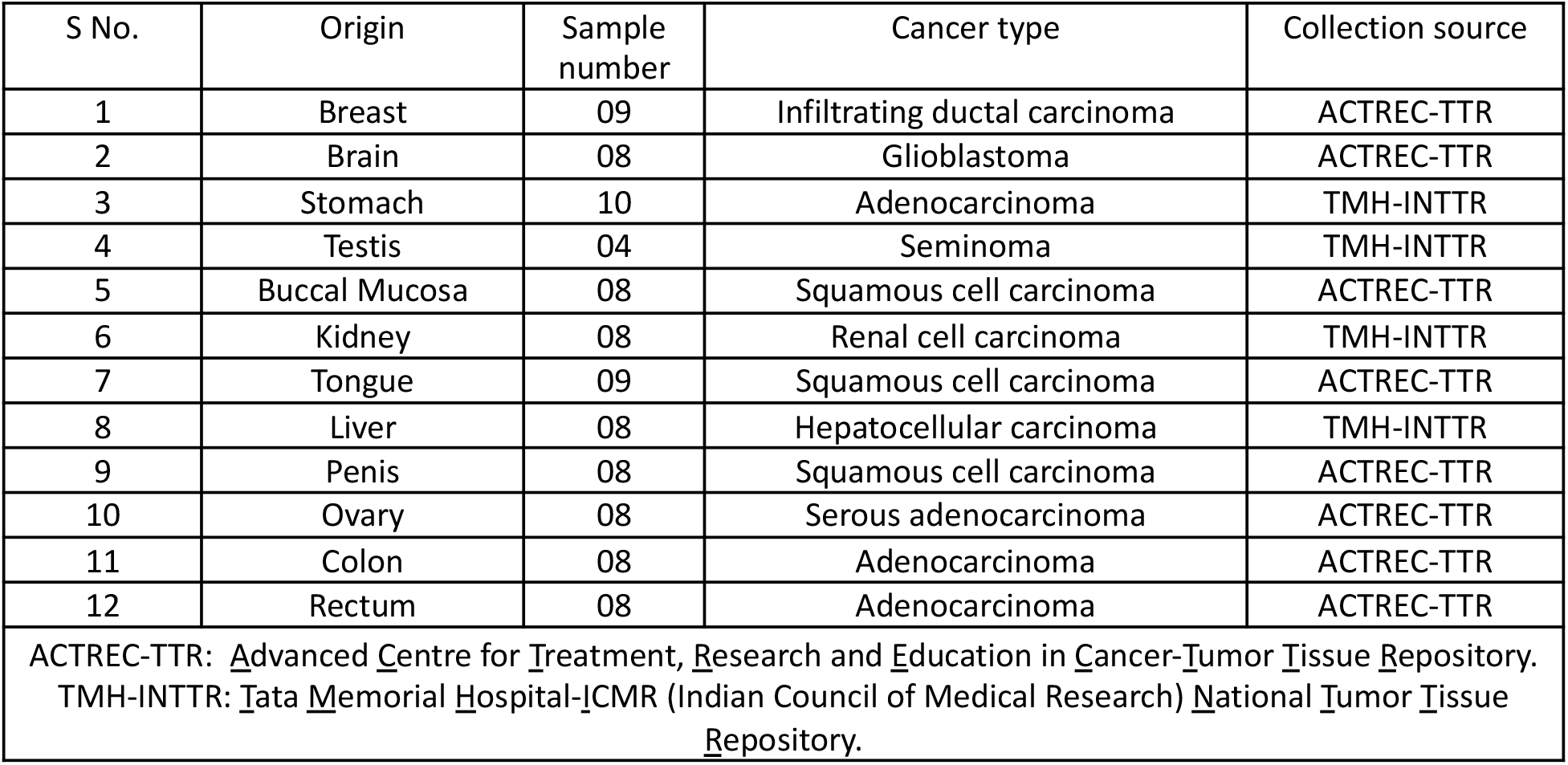
Details of human cancer tissue and cancer type: Figure representing twelve different tissue types along with their numbers, cancer type and collection source.

**Figure3-figure supplement1:**
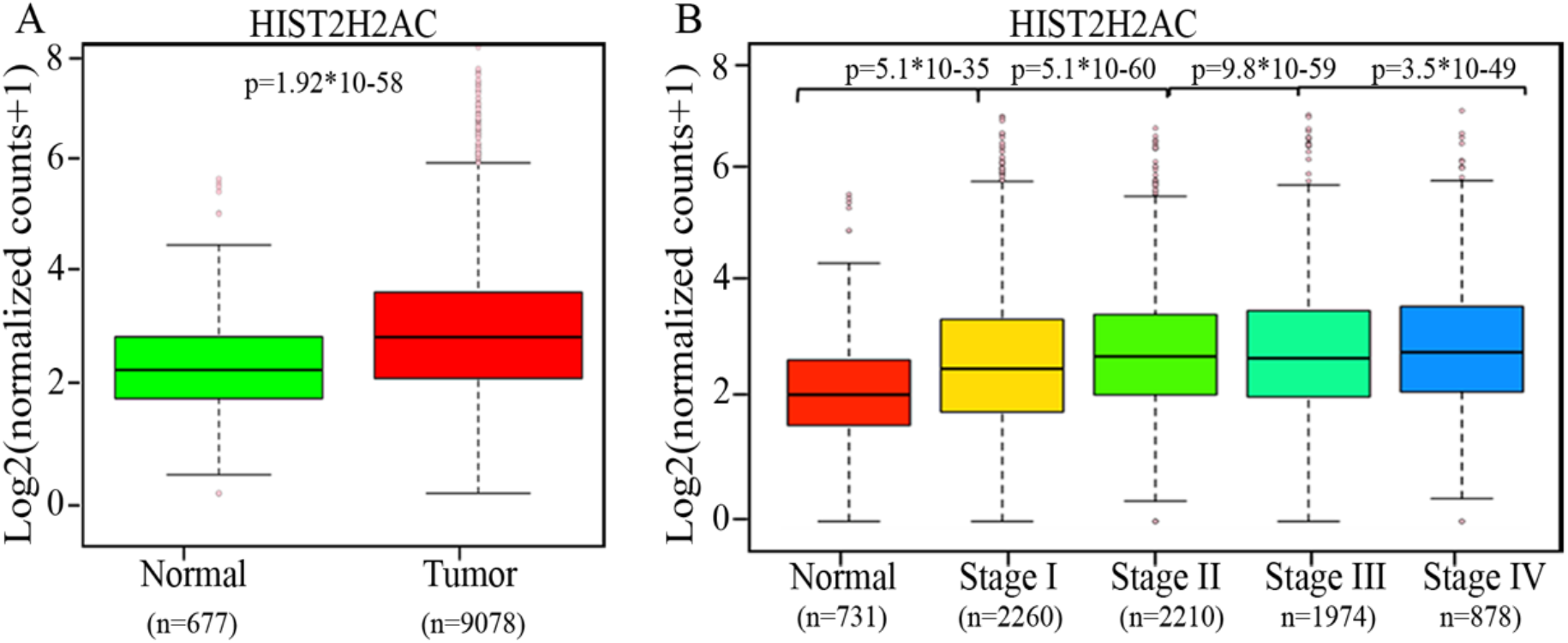
HIST2H2AC expression in TCGA Pancancer data: **(A)** Boxplot representation of normalized RSEM log2 (X+1) counts for HIST2H2AC expression in all normal and tumor samples (**B)** Boxplot representation of the expression of normalized RSEM log2 (X+1) counts for HIST2H2AC expression in all normal and different stages of tumor samples. p<0.05=*, p<0.01=** and p<0.001=***.

